# Latent cytomegalovirus disrupts NK cell responses to *P. falciparum* and impairs parasite control

**DOI:** 10.64898/2026.01.27.702133

**Authors:** Reena Mukhiya, Jessica R Loughland, Nick L Dooley, Zuleima Pava, Damian Oyong, Dean W Andrew, Julianne Hamelink, Kiana Berry, James S McCarthy, Bridget E Barber, J. Alejandro Lopez, Christian R Engwerda, Michelle J Boyle

## Abstract

**Background:** NK cells are major innate and adaptive responders to malaria, with multiple roles in protection. Function of NK cells is heterogeneous, underpinned by expression of a diversity of receptors. One driver of NK cell heterogeneity is latent CMV infection, which drives the expansion of memory-like NK cells. We have recently reported that latent CMV infection can negatively impact the adaptive immune response to malaria, but whether CMV-mediated changes to the NK cell compartment also impact innate responses to malaria is unknown.

**Methods:** We investigated the impact of latent CMV infection on NK cell response to the malaria parasite *Plasmodium falciparum in vitro*, and in CMV seronegative and seropositive individuals during controlled human malaria infection. We analysed NK cell activation, cytotoxicity and NK cell receptor expression. Additionally, we investigated the impact of CMV serostatus on cytokine production in response to TLR stimulation in the myeloid cell compartment. The impact of CMV and NK cell responses on parasite control and malaria symptoms was investigated.

**Results:** NK cells from CMV seropositive individuals had reduced responsiveness to *P. falciparum* parasites *in vitro* and had reduced activation during controlled human infection. Reduced activation was not restricted to NK subsets modulated by CMV but occurred across the entire NK cell compartment. Consistent with global NK cell attenuation, IL-12 production from myeloid cells, a response that supports NK cell activation on exposure to *P. falciparum* parasites, was lower in CMV infected individuals. Linking NK cell activation to clinical outcomes, NK cells expressing perforin were associated with parasite control in CMV seronegative individuals.

**Conclusion:** CMV infection modulates NK cell responses during malaria by disruption of IL-12, leading to reduced parasite control.

## Introduction

*Plasmodium falciparum* malaria remains a significant global health challenge, with an estimated 282 million cases and over 600 000 deaths in 2024 (World Health Organisation, 2025). Protective immunity to malaria is mediated by both innate and adaptive mechanisms that control parasites and limit immunopathology (Boyle et al., 2024). Natural Killer (NK) cells are key immune cells in both innate and adaptive responses to malaria and have multiple roles in protection (Goodier et al., 2019). NK cells are phenotypically heterogeneous, with this diversity and corresponding functionality influenced by multiple host and environmental factors (Horowitz et al., 2013; Rascle et al., 2023). One major driver of NK cell diversity is cytomegalovirus (CMV) infection which significantly modulates the NK cell compartment, driving expansion of ‘adaptive’ NK cells with hallmarks of memory including clonal expansion and modified responses to secondary challenge (Schlums et al., 2015; Goodier et al., 2018). We have recently shown that latent CMV infection is associated with reduced antibody development in malaria, both during infection and following vaccination (Mukhiya et al., 2024). However, whether latent CMV infection can also impact NK cell responses and the innate cellular response to malaria is unknown.

Human NK cells are defined based on CD16 and CD56 expression as CD16^-^CD56^bright^, CD16^+^ CD56^dim^ and CD16^+^CD56^neg^ cells and they express a vast array of activating and inhibiting receptors (Horowitz et al., 2013; Gonzalez et al., 2009; Mavilio et al., 2005; Ty et al., 2023). Recent studies have highlighted the key role of NK cells in malaria immunity, identifying NK cell receptors which can directly recognise *P. falciparum* parasites, and parasite-mediated NK cell inhibitor mechanisms (Sakoguchi et al., 2025; Saito et al., 2017). In response to parasites, NK cells rapidly produce IFNγ, a key cytokine for parasite control, along with cytotoxic molecules, a process that also requires NK cell activating cytokines IL12 and IL18 from innate cells (Artavanis-Tsakonas et al., 2003a; Artavanis-Tsakonas and Riley, 2002; Baratin et al., 2005; Newman et al., 2006). In mouse models, NK cell responses have been directly linked to parasite control and protection in some studies (Choudhury et al., 2000; Mohan et al., 1997). NK cells also have key roles in antibody-dependent cellular cytotoxicity (ADCC) via adaptive immune mechanisms which are associated with protection in children (Hart et al., 2019; Odera et al., 2023; Ty et al., 2023). The ability of NK cells to respond to malaria, through both innate and ADCC pathways, is heterogeneous in both malaria naïve and malaria exposed subjects. (Korbel et al., 2005; Artavanis-Tsakonas et al., 2003a; Artavanis-Tsakonas and Riley, 2002; Orago and Facer, 1991). This heterogeneity is in part mediated by underlying diversity in expression of NK cell receptors. For innate NK cell responses, a subpopulation of NK cells expressing the lectin-type receptor NKG2A is the predominant subset producing IFNγ in response to *P. falciparum* parasites *in vitro* (Artavanis-Tsakonas et al., 2003b). For NK adaptive immune responses, ADCC activity is highest in a specific subset of adaptive CD56^neg^ NK cells expressing increased LAG3 and LILRB1, with expansion of this subset also driven by malaria exposure (Ty et al., 2023; Hart et al., 2019).

One of the most significant drivers of heterogeneity of the NK cell compartment is CMV infection (Schlums et al., 2015). CMV is a ubiquitous beta-herpes virus which establishes an asymptomatic, life-long infection and has a global prevalence ranging from ∼50% in high income countries, to up to 100% in some areas of Africa where it is acquired early in life (Zuhair et al., 2019). Latent CMV drives expansion of adaptive CD56^dim^ CD57^+^ NKG2C^+^ NK cells which are characterized by reduced IFNγ and cytotoxic responses to exogenous IL12 and IL18 (Schlums et al., 2015; Gumá et al., 2004a). These CMV expanded adaptive NK cells, express lower frequencies of the Natural Cytotoxic Receptors (NCRs; for example, NKp30 and NKp46), and higher frequencies of inhibitory Killer-cell immunoglobulin-like receptors (KIRs) such as NKG2D (Béziat et al., 2013; Gumá et al., 2004b; Wu et al., 2013; Muntasell et al., 2013). These changes have important consequences for responsiveness of NK cells to other pathogens and stimuli. For example, both IFNγ production and degranulation are reduced in CD57^+^NKG2C^+^ NK cells when Peripheral Blood Mononuclear Cells (PBMCs) from CMV seropositive UK adults are stimulated with pertussis or H1N1 influenza vaccine antigens (Nielsen et al., 2015). Here we investigate the impact of CMV on NK cell responsiveness to *P. falciparum* parasites, with a focus on innate cell responses in malaria naïve individuals and explore the consequences of changes to parasite control.

## Results

### The transcriptional response of NK cells to *P. falciparum* is modulated by latent CMV infection

To dissect the impact of latent CMV infection on NK cell responses to malaria, we first assessed the transcriptional profile of NK cells from CMV seronegative and CMV seropositive healthy, malaria naïve individuals following *in vitro* stimulation with *P. falciparum* parasite infected red blood cells (pRBCs). PBMCs were cultured for 24 hours with parasites or media, and then NK cells isolated with negative bead selection for sequencing (CMV seronegative n=6, age range 29-36, 50% male and CMV seropositive n=6, age range 26-45, 50% male) (Supplementary Figure S1A/B). NK cells from CMV seronegative and CMV seropositive donors were transcriptionally distinct prior to and following stimulation (Figure 1A). Differentially expressed genes (DEGs) with CMV serostatus (negative compared to positive), parasite stimulation (unstimulated compared to stimulated), and genes which responded in a CMV specific manner (significant for CMV serostatus and stimulation, and/or with a significant interaction term between CMV and stimulation) were identified using *glmmSeq* (Lewis et al., 2022), which fits a negative binomial mixed-effects model at the individual gene level. Using this approach, 323 DEGs were identified that were modulated by CMV serostatus, consistent with the reported modulations of NK cells in CMV infection (Gumá et al., 2004b; Lopez-Vergès et al., 2011a; Béziat et al., 2012) (Supplementary Figure S1C, Supplementary Table S1). In unstimulated cells, DEGs upregulated in CMV seropositive individuals included known markers of CMV-mediated NK cell changes, including *B3GAT1 (*encoding CD57) (Lopez-Vergès et al., 2011b), and *LILRB1* (Cadena-Mota et al., 2018) (Supplementary Figure S1E). While *KLRC2* (encoding NKG2C) was not identified, this gene was significantly higher in CMV seropositive individuals in unadjusted analysis (p = 0.004, q (adjusted)=0.11) (Supplementary Table S1).

**Figure 1:**
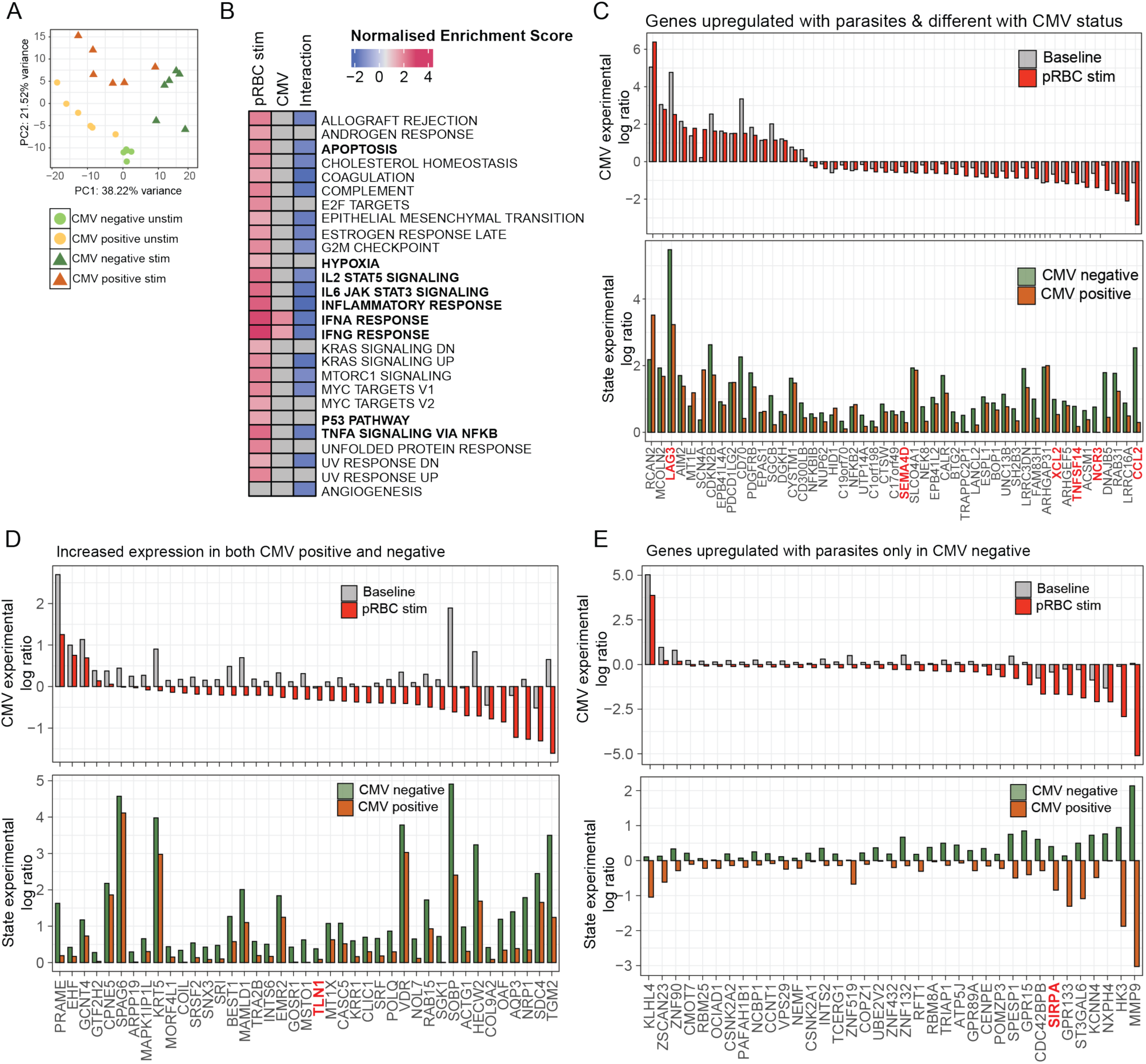
Transcriptional profiling of NK cells after stimulation with P. falciparum in CMV seronegative and seropositive individuals. **(A)** NK cells from CMV seronegative (n=6) and CMV seropositive (n=6) individuals were analysed by RNAseq ex vivo and following stimulation with P. falciparum infected RBC (pRBC). Principal component analysis (PCA) of DEGs identified by glmmSeq, with individuals’ CMV and stimulation status indicated. **(B)** GSEA pathway analysis of all DEGs identified as significantly modulated by parasite stimulation (pRBC), CMV serostatus, and either both or an interaction term. Normalised enrichment score for hallmark pathways shown. **(C)** DEGs that were significant for CMV status and were upregulated following stimulation, but without interaction term. **(D/E)** DEGs that had a significant interaction term between CMV serostatus and stimulation, which were upregulated in both CMV seronegative and CMV seropositive (**D**), or only upregulated in CMV seronegative individuals (**E)**. For (**C-E)** Top panels are the CMV serostatus log ratio at baseline (grey) or following stimulation (red). A positive log ratio indicates genes which were significantly higher in CMV positive individuals. Bottom panels are State log ratio in CMV seronegative (green) and CMV seropositive (orange) individuals. A positive log ratio indicates genes that were upregulated following stimulation with pRBCs. See also Supplementary Figure 1 and Supplementary Table S1.

In response to parasite stimulation, 9578 DEGs were identified, consistent with a robust response of NK cells to parasites (Supplementary Figure 1C, Supplementary Table S1). Of these genes, 259 (2.7%) also differed significantly by CMV serostatus (indicating that the expression differed by CMV status before and after stimulation, 148 DEGs) and/or had a significant interaction with stimulation (indicating that the response to parasites were different based on CMV serostatus, 111 DEGs). Gene set enrichment analysis (GSEA) of DEGs identified upregulation of major inflammatory pathways including in response to parasite stimulation (Figure 1B). However, for DEGs modified by CMV only IFNA and IFNG response pathways were identified. For DEGs that were identified for both parasite stimulation and CMV serostatus, or had an interaction term, GSEA showed down regulation of hallmark signatures including inflammatory pathways that were otherwise upregulated in response to parasites, suggesting that NK cells from CMV seropositive individuals had an attenuated response to parasite stimulation. Indeed, analysis of individual DEGs that were identified for both parasite stimulation and CMV, or had a significant interaction term, suggested important functional differences (Figure 1C/D/E). For example, genes upregulated in response to parasites included *LAG3,* expression of which was higher in CMV seropositive individuals both before and after stimulation (Figure 1C). The inhibitory receptor LAG3 has recently been shown to inhibit NK cell IFNγ production and proliferation both in healthy and HIV infected individuals (Ge et al., 2025). In contrast, genes *CCL2* (which is essential for monocyte recruitment) and *NCR3 (*encoding the NK cell activating receptor NKp30) were higher in CMV seronegative individuals, both before and after parasite simulation (Figure 1C). Furthermore, *SEMA4D* (CD100), reported to enhance NK cell killing (He et al., 2017); *TNFSF14,* which has been shown to increase with NK cell activation and induce DC maturation (Holmes et al., 2014); and *XCL2,* a key NK cell chemoattractant that recruits DCs (Böttcher et al., 2018), were all higher in CMV seronegative individuals. Of DEGs with a significant interaction term between stimulation and CMV serostatus (indicating a significantly different response to parasites between the groups), 41 DEGs were significantly less activated in CMV seropositive individuals and/or significantly lower in CMV seropositive individuals following parasite stimulation, and a further 36 DEGs were only upregulated in CMV seronegative individuals (Figure 1D/E). Amongst these DEGs were *TLN1* (encoding talin-1), essential for NK cell cytotoxicity by mediating adhesion and cell polarization (Mace et al., 2009), and *SIRPA* (encoding SIPRα), which is upregulated in NK cells after cytokine activation (Deuse et al., 2021). Taken together these DEGs suggest increased NK cell functional responses to *P. falciparum* parasite stimulation in CMV seronegative individuals.

### NK cell activation during a first malaria infection is reduced in CMV seropositive individuals

To assess if CMV serostatus modulated NK cell responses to *P. falciparum* parasites *in vivo,* we analysed responses in individuals during controlled human malaria infection (CHMI) (CMV seronegative n= 10, 26 years median (IQR 20-31), male 90%, CMV seropositive n=11, 25 years median (IQR 21-29), male 72.7 %) (Mukhiya et al., 2024; Chan et al., 2020). NK cells were analysed based on CD56 expression to identify CD56^bright^, CD56^dim^ and CD56^neg^ subsets (Supplementary Figure S2A). Overall, there were limited differences between CMV seronegative and seropositive donors for total frequencies of NK cells. Further, the proportions CD56^bright^, CD56^dim^ cells were similar before and during CHMI (Supplementary Figure S2B). Following *P. falciparum* infection, there was a significant increase in CD56^bright^ NK cells and a decrease in CD56^neg^ NK cells in both CMV seronegative and CMV seropositive individuals (Supplementary Figure S2B-C).

To quantify NK cell responsiveness during malaria, activation and cytotoxicity was quantified as the median signal intensity of CD38, NKp30, Perforin, Granulysin and GranyzmeB on each NK subset, and analysed with linear mixed effects models (Supplementary Figure S2A). For CD56^dim^ NK cells, activation markers CD38 and NKp30 significantly increased expression in both CMV seronegative and seropositive individuals, with the highest expression at day 15 post infection (Figure 2A). However, the magnitude of the increase was significantly higher for CD38 in CMV seronegative individuals (mixed model interaction term p=0.007) (Figure 2A). For cytotoxic markers Perforin, Granulysin and GranyzmeB, increased expression was only detected in CMV seronegative individuals following CHMI (Figure 2B). For CD56^bright^ cells, the increased activation and cytotoxic activity in CMV seronegative individuals was even more striking, with a significantly higher increase of all markers at day 15 (Supplementary Figure S2D). Similarly, for CD56^neg^ NK cells, an increased response for CD38, Granulysin and Granzyme B was also observed in CMV seronegative individuals (Supplementary Figure S2E). Together these data suggest that CMV infection has a negative impact on malaria-induced activation of NK cells across all NK cell subsets.

**Figure 2:**
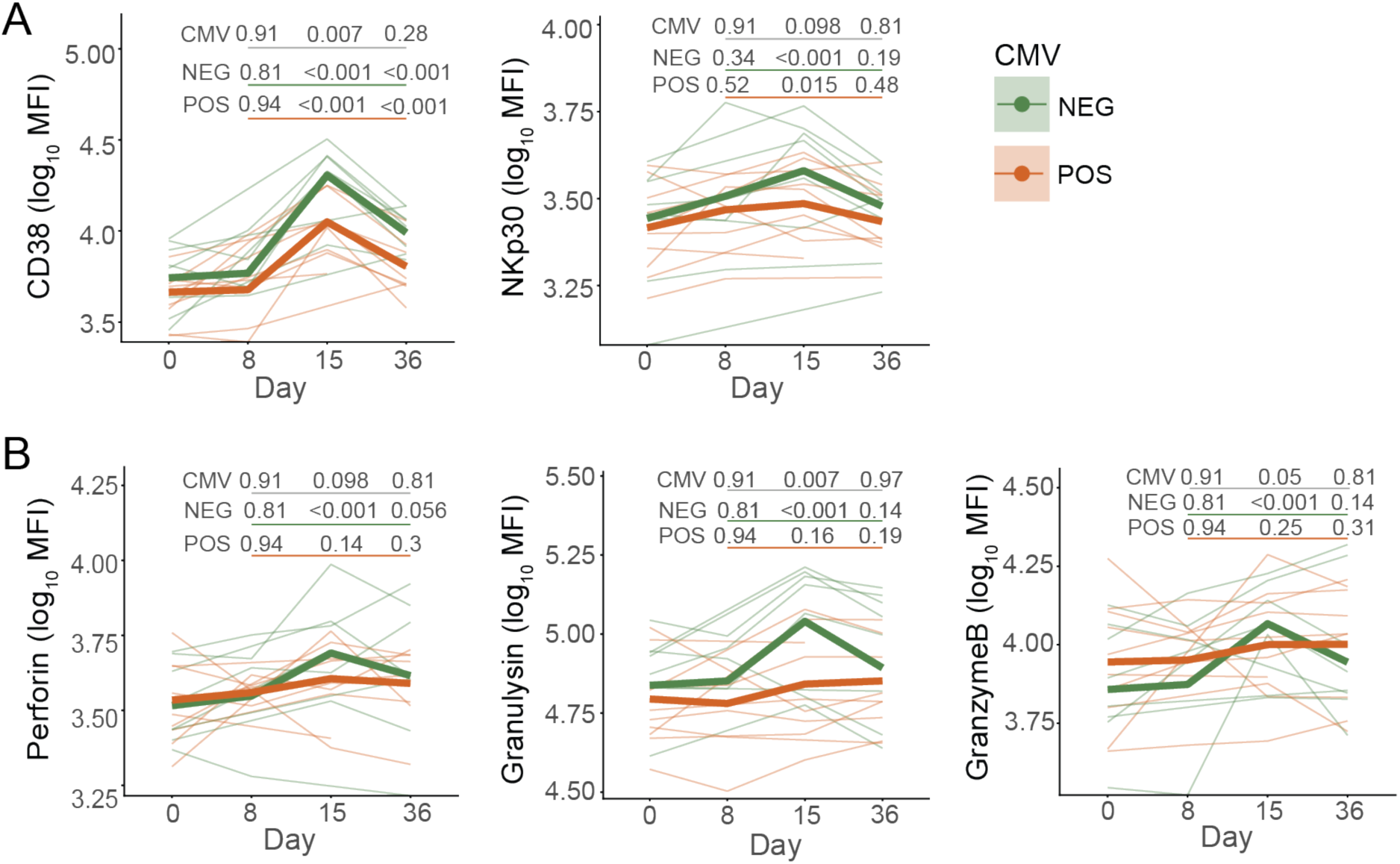
CD56^dim^ NK cell activation during controlled human malaria infection in CMV seronegative and seropositive individuals. CD56^dim^ NK cell activation was analysed by quantifying median fluorescence intensity (MFI) of **(A)** activation markers CD38 and NKp30, and **(B)** cytotoxic markers Perforin, Granzyme B and Granulysin in CMV seronegative (n=8) and CMV seropositive individuals (n=9). Data are log_10_ MFIs of markers with thin lines representing individual data coloured by CMV serostatus, and bold lines representing the mean of the predicted values from the fitted models for each group. P values are from linear mixed effect models. CMV is p values for the interaction term between each timepoint (compared to day 0) and CMV serostatus (underlined in grey). NEG/POS are P values for the comparison between day 0 and each subsequent timepoint for CMV seronegative individuals (NEG, underlined in green) and CMV seropositive individuals (POS, underlined in orange) which were determined from contrasts. See also Supplementary Figure S2.

### CMV associated changes to the NK cell compartment are maintained during CHMI

NK cell phenotypes are modulated by CMV, which induces expansion of adaptive CD56^dim^ CD57^+^/NKG2C^+^ NK cells, resulting in a proportional decrease of NKG2A^+^ NK cells within CMV seropositive individuals (Schlums et al., 2015). Because NKG2A^+^ NK cells preferentially respond to *P. falciparum in vitro* (Artavanis-Tsakonas et al., 2003a), we hypothesised that reduced activation of NK cells during malaria may be due to reduced frequencies of this subset. To investigate this hypothesis, the NK cell population was analysed based on expression of NKG2A, NKG2C, CD57, together with CD56. Clustering of NK cells based on these markers identified nine NK cell clusters which included two CD56^bright^ subsets, one co-expressing NKG2A, and the other expressing both NKG2A and NKG2C; six CD56^dim^ subsets, including a CD57^+^NKG2C^+^ subset, and a NKG2A^+^ subset; and a small cluster of CD56^neg^ cells (Figure 3A). Distribution of subsets was heterogenous at the individual level (Figure 3B). However, consistent with prior studies, there was a significantly increased proportion of CD57^+^NKG2C^+^CD56^dim^ NK cells (p=0.02), and a decreased proportion of NKG2A^+^CD56^bright^ (p=0.05) and NKG2A^+^CD56^dim^ (p=0.11) cells in CMV seropositive individuals prior to malaria (Figure 3C). For CD57^+^NKG2C^+^CD56^dim^ cells, these CMV-associated differences were maintained over the course of infection (Figure 3D/E). For NKG2A^+^ subsets, CMV-associated differences were only maintained for NKG2A^+^CD56^dim^ and not NKG2A^+^CD56^bright^ cells, the latter of which expanded in both CMV seronegative and CMV seropositive individuals during CHMI (Figure 3D/E).

**Figure 3:**
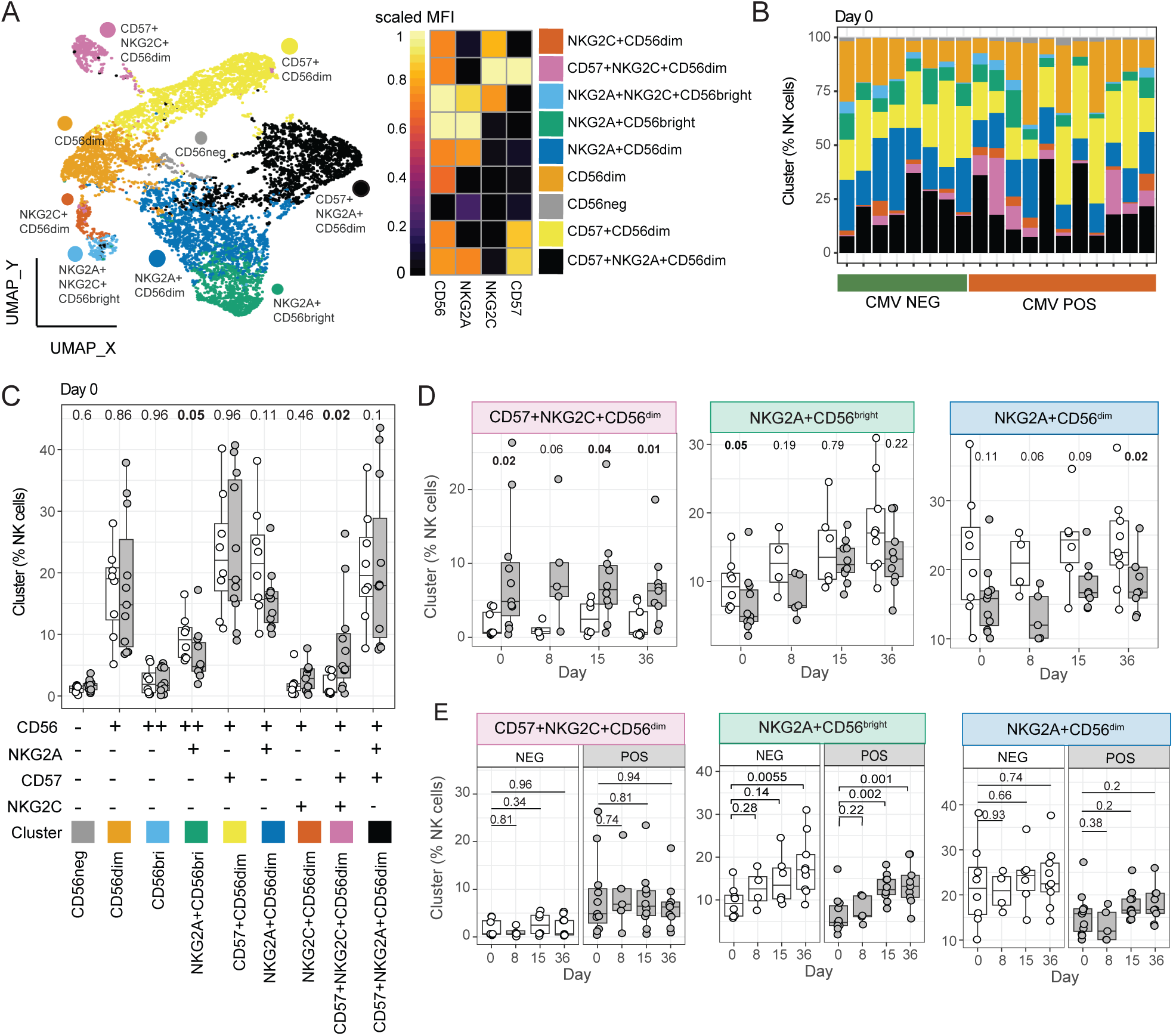
NK cell subset distribution in CMV seronegative and seropositive individuals during CHMI. **(A)** NK cell phenotypes were classified based on expression of CD57, NKG2A, NKG2C and CD56. UMAP projection of NK cell clusters, and heatmap of cluster maker expression on each subset. **(B)** Percentage of each identified NK cell cluster in each CMV seronegative (n=8) and CMV seropositive (n=9) individual at day 0, prior to malaria inoculation. **C)** Proportion of NK cell subsets between CMV seronegative and CMV seropositive individuals. Marker expression of each cluster is indicated. **(D/E)** CD57+NKG2C+CD56^dim^, NKG2A+CD56^bright^ and NKG2A+CD56^dim^cells comparing CMV seronegative and CMV seropositive individuals at each CHMI timepoint **(D)** and comparing day 0 to each subsequent timepoint (**E)**. For **(C-E)** data are Tukey boxplots with the median, 25^th^ and 75^th^ percentiles. The upper and lower hinges extend to the largest and smallest values, respectively but not further than 1.5X IQR from the hinge. Individual data are shown as points. For **C/D** p is Mann-Whitney U test. For **E** p is Wilcoxon signed-rank test.

### CMV seropositive individuals have reduced activation and cytotoxic response during malaria across multiple NK cell subsets

To assess if CMV-associated changes to the NK cell compartment underpinned the impaired NK cell responsiveness during malaria, we analysed activation (CD38 and NKp30) and cytotoxicity (Perforin, Granulysin, and Granzyme B) across all identified subsets. We focused on day 15 after CHMI which was the peak of NK cell activation (Figure 2) and focused on the NK cell subsets that were influenced by CMV serostatus (Figure 3). In contrast to *in vitro* reports of higher responsiveness to *P. falciparum* in NKG2A^+^ NK cells (Artavanis-Tsakonas et al., 2003a), there was robust activation of CD57^+^NKG2C^+^CD56^dim^, and NKG2A^+^CD56^bright^ and NKG2A^+^CD56^dim^ subsets, all of which had increased CD38 expression at day 15 in both CMV seronegative and CMV seropositive individuals (Figure 4A). At day 15, CD38 expression was higher in CMV seronegative individuals for all subsets (Figure 4A). CHMI mediated increases of NKp30 were less robust and did not reach statistical significance in either CMV seronegative or CMV seropositive groups for all subsets (Figure 4B). For cytotoxic markers, there was also evidence for stronger activation at day 15 of CHMI in CMV seronegative individuals across subsets, with significantly increased Perforin expression in both NKG2A^+^ subsets (Figure 4C), increased Granulysin expression across all subsets (Figure 4D), and increased Granzyme B expression in NKG2A^+^CD56^bright^ cells (Figure 4E). In contrast, for CMV seropositive individuals, malaria only drove increased cytotoxic markers for Granulysin in NKG2A^+^CD56^bright^ cells (Figure 4C-E). This pattern of more robust activation and larger increases in cytotoxic markers in CMV seronegative subjects was seen in all other NK cell subsets (Supplementary Figure 3A-E). For CD38, while expression increased at day 15 for most subsets (all except CD56^neg^ cells) regardless of CMV infection status, expression was significantly higher in CMV seronegative individuals (Supplementary Figure 3A). Further, for NKp30, there was increased expression at day 15 after CHMI in 4/6 additional NK cell subsets, but no induction detected in CMV seropositive individuals (Supplementary Figure 3B). Similarly, Perforin expression increased in 5/6 subsets in CMV seronegative individuals and only in 1/6 for CMV seropositive; Granulysin increased in 4/6 subsets in CMV seronegative subjects but did not increase in any CMV seropositive individuals (Supplementary Figure 3C-D). For Granzyme B, responses were more varied; increased expression only reached statistical significance for a single subset amongst the CMV seropositive individuals (Supplementary Figure 3E). Taken together, these data indicate that CHMI induces robust activation across multiple NK cell subsets, and that increases in cytotoxic marker expression are consistently more robust in CMV seronegative individuals.

**Figure 4:**
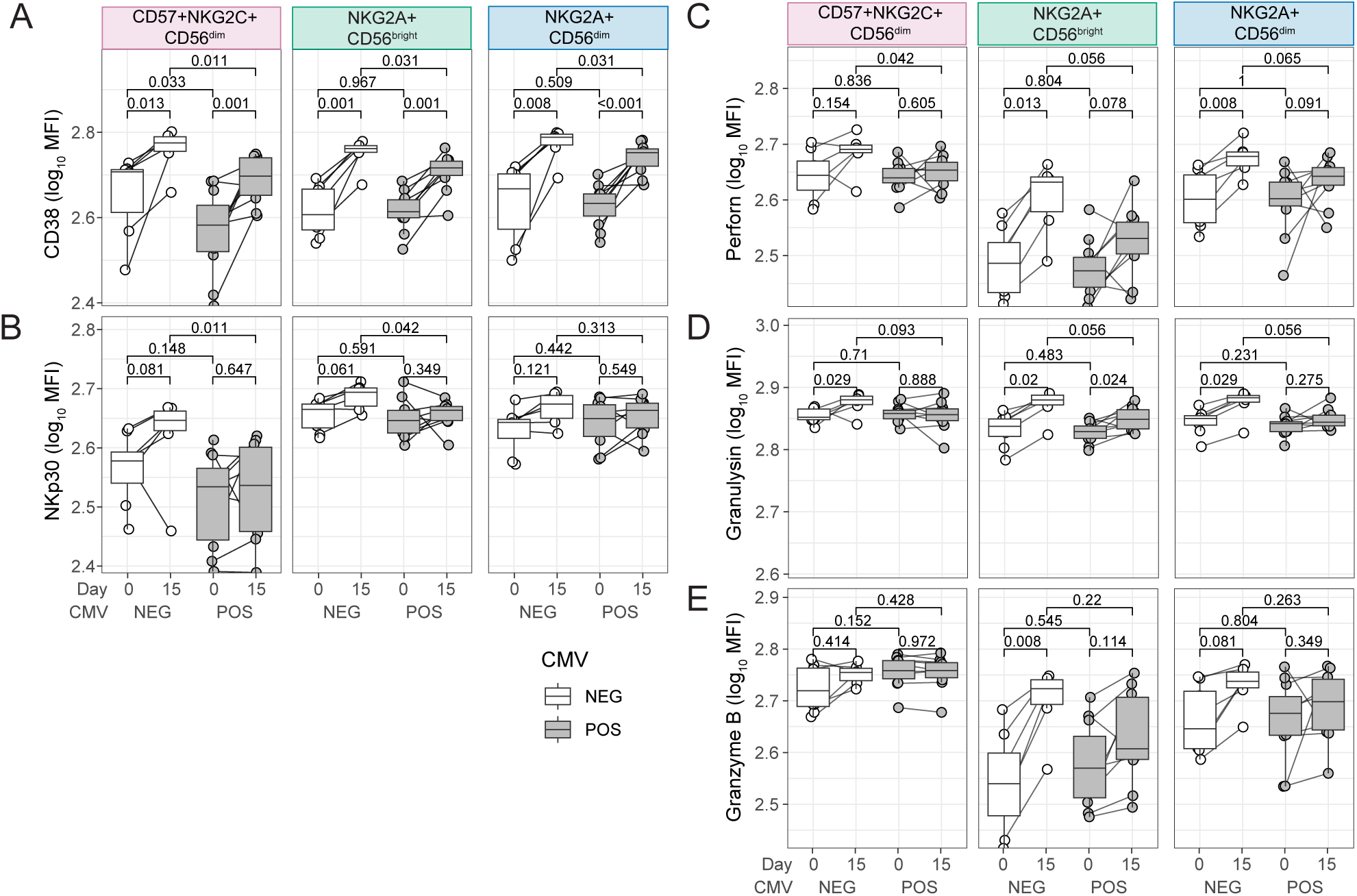
Activation and cytotoxicity in CMV modulated subsets during CHMI. Activation (**A** - CD38 and **B** - NKp30) and cytotoxic (**C** - Perforin, **D** - Granulysin, **E** – Granzyme B) markers were quantified (median fluorescence intensity, MFI) in NK cell subsets that were modulated by CMV infection, CD57+NKG2C+CD56^dim^, NKG2A+CD56^bright^ and NKG2A+CD56^dim^ subsets. Expression was compared between day 0 and day 15 in CMV seronegative (n = 8) and CMV seropositive individuals (n = 9), and expression at day 0 or at day 15 was compared between groups. Data are Tukey boxplots with the median, 25^th^ and 75^th^ percentiles. The upper and lower hinges extend to the largest and smallest values, respectively but not further than 1.5X IQR from the hinge. Individual data are shown as points. For comparisons between groups p is Mann-Whitney U test. For comparisons within groups between days P is Wilcoxon signed-rank test. See also Supplementary Figure S3.

### Monocyte IL12 production in response to TLR4 stimulation is higher in CMV negative individuals

In previous studies, it has been shown that together with parasite contact, the cytokines IL12 and IL18 are essential for NK cell activation *in vitro* by *P. falciparum* (Artavanis-Tsakonas et al., 2003a; Artavanis-Tsakonas and Riley, 2002; Baratin et al., 2005; Newman et al., 2006). Because our data suggested that NK cell activation and toxicity was disrupted across multiple subsets during CHMI in CMV seropositive individuals, we hypothesized that CMV infection may reduce the cytokine milieu from innate cells that supports NK cell activation. To test this hypothesis, we stimulated whole blood with innate cell agonists Pam3Csk4/HKLM, LPS and CL075, which activate via TLR1/2, TLR4 and TLR7/8 respectively (CMV seronegative n= 12, age range 18-53 years, male 50%, CMV seropositive n=17, age range 19-55 years, male 58.8%). These TLR pathways have been shown to be activated by *P. falciparum* including: TLR2 and TLR4 recognition of Glycosylphosphatidylinositol (GPI) anchors (Krishnegowda et al., 2005), TLR4 binding hemozoin bound with parasite proteins (Barrera et al., 2011), and TLR8 responding to *Plasmodium* RNA (Coch et al., 2019). Following stimulation, IL12, TNF and IL6 were quantified by intracellular staining in classical and non-classical monocytes and dendritic cells (DCs) (Supplementary Figure 4A/B). In both classical monocytes and dendritic cells, significantly lower IL12 production was measured in response to TLR4 stimulation in CMV seropositive individuals (Figure 5A). There was also a significant reduced IL12 production in CMV seropositive individuals in response to TLR1/2 stimulation in non-classical monocytes (Figure 5A). In contrast, TNF production was higher in CMV seropositive individuals after TLR4 stimulation in non-classical monocytes and TLR1/2 stimulation in dendritic cells (Figure 5B). Additionally, levels of IL6 were higher in CMV seropositive individuals after TLR1/2 and TLR4 stimulation in non-classical monocytes, and TLR1/2 stimulation for dendritic cells (Figure 5C).

**Figure 5:**
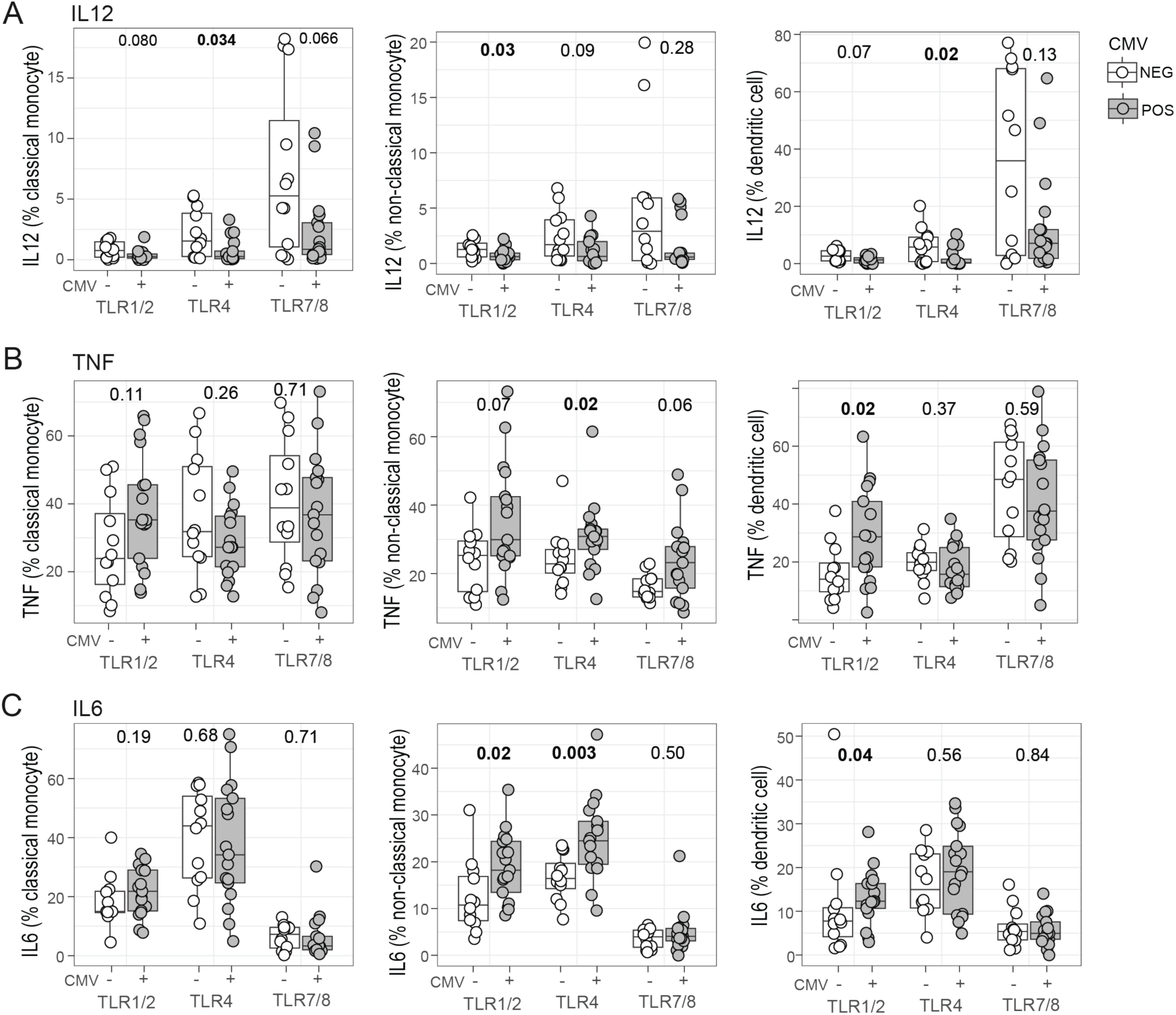
Cytokine production following TLR stimulation in myeloid cells in CMV seronegative and seropositive individuals. Whole blood from healthy CMV seronegative (n=12) and CMV seropositive (n=17) donors was stimulated with TLR agonists Pam3Csk4/HKLM for TLR1/2, LPS for TLR4 and CL075 for TLR7/8 and IL12 **(A**), TNF **(B)** and IL6 **(C)** quantified in classical monocytes, non-classical monocytes and dendritic cells. See also Supplementary Figure S4.

### Parasite multiplication rate is higher in CMV seropositive individuals and associated with NK cell activation

To assess the clinical relevance of CMV-mediated changes to NK cells and other innate cell responses to malaria, we analysed associations between CMV serostatus and early parasite control and inflammation in 40 individuals during CHMI (Supplementary Table S2). In this cohort, we recently reported that CMV negatively impacts antibody induction but does not associate with the parasite burden observed over the total infection period (before and after treatment) (Mukhiya et al., 2024). However, total parasite burden is influenced by both parasite growth and drug efficacy, therefore we assessed the association between CMV and parasite multiplication rate (PMR) prior to treatment. PMR was significantly higher in CMV seropositive individuals (Figure 6A). In contrast, the combined malaria clinical score, an indicator of inflammation (Webster et al., 2025), was significantly lower in CMV seropositive individuals, particularly early in infection (Figure 6B).

**Figure 6:**
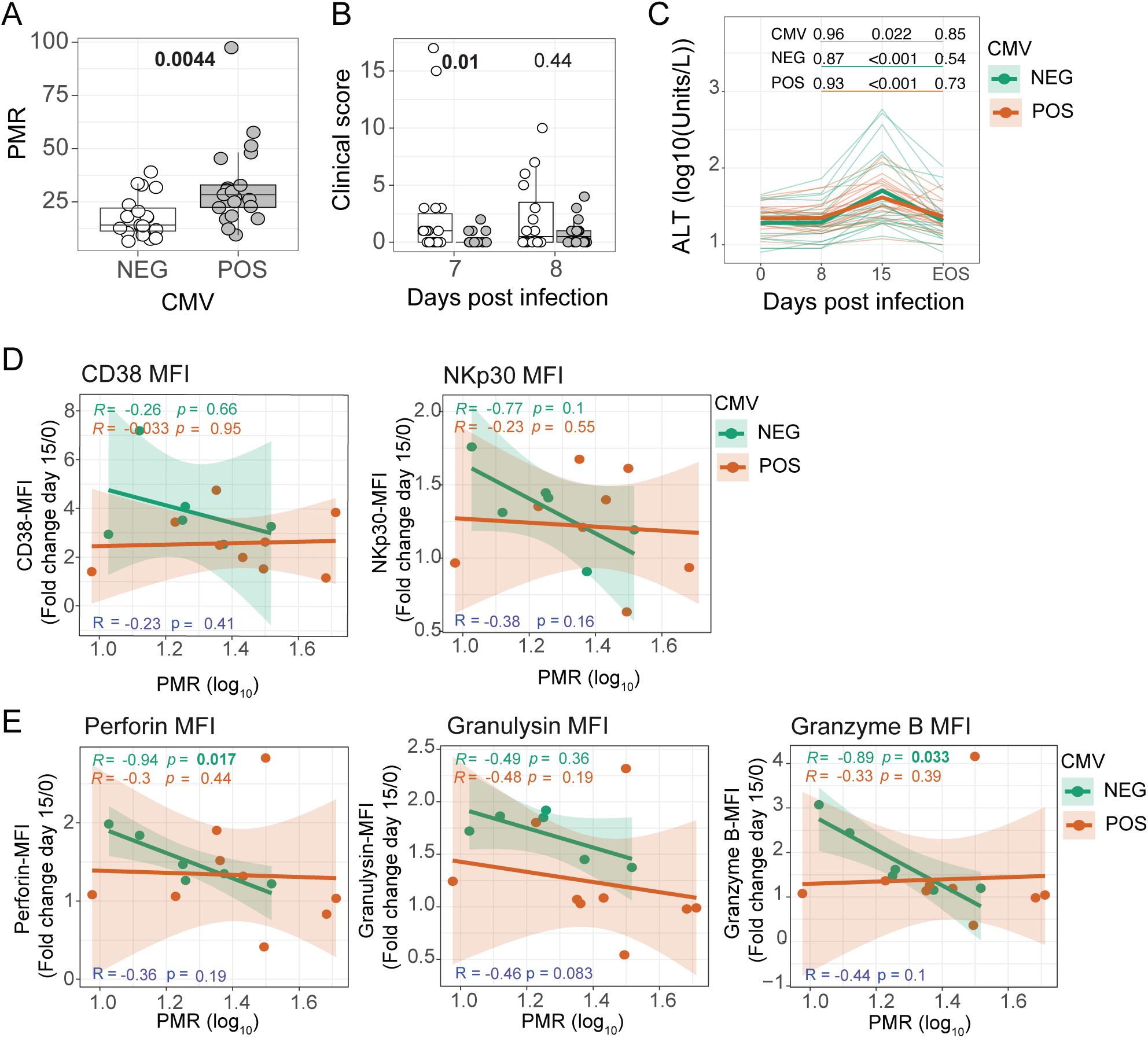
Parasite multiplication rate, clinical parameters and NK cell associations. Associations between CMV serostatus and parasite control and indicators of inflammation were investigated in a cohort of individuals in CHMI (CMV seronegative n= 19, CMV seropositive n=21). **A**) Parasite multiplication rate (PMR) was compared between CMV seronegative and seropositive individuals. **B)** Clinical score prior to treatment between CMV seropositive and seronegative. For **A-B** data are Tukey boxplots with the median, 25^th^ and 75^th^ percentiles. The upper and lower hinges extend to the largest and smallest values, respectively but not further than 1.5X IQR from the hinge. Individual data are shown as points. For comparisons between groups p is Mann-Whitney U test. For **A/B** p is wilcoxon signed-rankest test. **C)** Alanine transaminase at day 0, 8, 15, EOS. Data are log_10_ ALT with thin lines representing individual data coloured by CMV serostatus, and bold lines representing the mean of the predicted values from the fitted models for each group. P values are from linear mixed effect models. CMV is p values for the interaction term between each timepoint (compared to day 0) and CMV serostatus (underlined in grey). NEG/POS are P values for the comparison between day 0 and each subsequent timepoint for CMV seronegative individuals (NEG, underlined in green) and CMV seropositive individuals (POS, underlined in orange) which were determined from contrasts. **D/E)** Fold change increase in expression of activation markers CD38 and NKp30 **(D)**, or cytotoxic markers Perforin, Granulysin, or Granzyme B **(E)** on CD56^dim^ NK cells was calculated for day 15 during CHMI and correlated with parasite multiplication rate (PMR) in a subset of individuals. CMV seronegative individuals are green (n=8) and CMV seropositive individuals are shown in orange (n=9). Spearman’s Rho and p are shown for CMV seronegative (green), CMV seropositive (orange) or all individuals (purple).

Consistent with this, the malaria induced increase of alanine transaminase (ALT), a marker of tissue damage, was significantly greater in CMV seronegative individuals (Figure 6C). To link these CMV-associated changes to parasite control to reduced activation of NK cells in CMV seropositive subjects, we assessed the association between activation of CD56^dim^ NK cells and PMR. There was no association between the magnitude of activation (fold increase of CD38 or NKp30 at day 15 compared to baseline) and PMR in either CMV seronegative, CMV seropositive, nor total individuals (Figure 6D). However, there was a strong and significant negative correlation between the increase in cytotoxic markers Perforin and Granzyme B and PMR in CMV seronegative individuals (Figure 6E). This relationship was not seen in CMV seropositive individuals, or when CMV serostatus was not considered. Together, these data indicate CMV infection negatively reduces the capacity of NK cells to respond to *P. falciparum* parasites, reducing parasite control and malaria symptoms.

## Discussion

NK cells have key roles in protective immunity, participating in both innate and adaptive responses to pathogens, including malaria. Here we show that the innate NK cell responses to malaria are significantly modified by latent CMV. CMV seropositive individuals have reduced transcriptional responsiveness to *P. falciparum* parasite stimulation *in vitro* and reduced NK cell activation and cytotoxicity to malaria *P. falciparum* infection *in vivo*. This attenuated responsiveness to parasites was seen in all NK cell subsets and was not limited to NK cell subsets expanded by CMV infection, suggestive of a systemic impact of CMV, possibly within myeloid cells. Of clinical relevance, the PMR was greater in CMV seropositive individuals, which was associated with reduced NK cell cytotoxic responsiveness. Together, data identify an important influence of CMV on NK cell function and response to *P. falciparum* infection and malaria.

Heterogeneity in human immune responses to pathogens is well recognised, and in malaria can range from low grade, controlled and asymptomatic, to high parasitemia and death. Underpinning these heterogeneous outcomes is the intersection of host and parasite factors, including the expression of parasite virulence genes (Kessler et al., 2017; Jespersen et al., 2016), host genetics (Band et al., 2013), age (Loughland et al., 2025; Baird, 2017; Dondorp et al., 2008; Barber et al., 2017) and prior exposure and acquired immunity (Doolan et al., 2009). Our findings highlight how individual pathogen exposure histories influences immune activation and disease outcome. Specifically, we show that latent CMV modulates host responses to *P. falciparum*, particularly within the NK cell compartment. Latent CMV infection drives the expansion of ‘adaptive’ CD56^dim^ NK cells, expressing both CD57 and NKG2C, and proportional decreases in NKG2A (Schlums et al., 2015). While we also confirmed these CMV-associated changes within NK cells within our cohorts, these changes did not appear to directly underpin reduced responsiveness to *P. falciparum*. Indeed, despite *in vitro* studies suggesting NKG2A is important for malaria responsiveness (Artavanis-Tsakonas et al., 2003a), here we observed NK cell activation across all subsets during CHMI, which was globally attenuated in CMV seropositive individuals. Consistent with this global attenuation, IL12 production from myeloid cells was reduced, a key supporting cytokine for NK cell activation in response to parasites. Thus, data suggest that the reduced responsiveness of NK cells to *P. falciparum* is not cell intrinsic, but instead due to CMV-mediated changes to other cell compartments.

Alongside major remodelling of the NK cell compartment, latent CMV drives other changes across the immune landscape, with a systems analysis reporting that half of all measured immune parameters were impacted by CMV serostatus (Brodin et al., 2015). Consistent with these previous findings, we show the cytokine production in response to different TLR agonists and transcriptional responses to *P. falciparum* were modulated by CMV serostatus. Specifically, the production of the NK-activating cytokine IL12 was lower in CMV seropositive individuals. *In vitro* studies have shown that CMV viral homology to IL10 (cmvIL10) can inhibit IL12 secretion induced by TLR4 (LPS) in DCs (Chang et al., 2004). However, in the same *in vitro* system, cmvIL10 also reduced IL6 and TNF, in contrast to data presented here where these cytokines were elevated in CMV seropositive individuals. While CMV DNA is rarely detected in circulating cells (Parry et al., 2016; Jackson et al., 2017), it has been detected in bone marrow CD34^+^ hemopoietic progenitor cells from healthy CMV seropositive individuals (Minton et al., 1994). Analogous to innate cell training, it is possible that latent infection within the hemopoietic compartment imprints myeloid cells to alter their responsiveness to pattern recognition signals (Netea et al., 2020). Thus, further studies are required to better understand how latent CMV infection within stem cells or elsewhere modulates the myeloid cell compartment and the impact of these changes to NK cell responsiveness to *P. falciparum* and other stimuli.

NK cells have key roles in protection from malaria, implicated as both innate cell responders and adaptative cells that control parasite growth via ADCC (Boyle et al., 2024). This study solidifies the importance of the NK cell response in early parasite control, identifying that NK cell cytotoxicity is associated with PMR during first *P. falciparum* infection in CMV seronegative individuals. Additionally, we have previously reported that latent CMV infection is associated with reduced antibody induction in the same controlled human malaria infection study, highlighting that CMV has negative impacts on malaria immunity in both innate and adaptive responses (Mukhiya et al., 2024). Given that CMV is acquired very early in life in malaria endemic areas, our data highlight the importance of better understanding the immunomodulatory impact of CMV in children living in malaria endemic areas. Indeed, while data here suggest that CMV may reduce parasite control, clinical symptoms were also reduced in CMV seropositive individuals. The impact of CMV on the severity of natural infection is unknown. While reduced capacity to control parasites may indicate increased risk of hyper-parasitemia and severe disease, paradoxically, the reduced inflammatory response may be protective from immunopathogenesis. Indeed, while CHMI studies are powerful approaches to investigate immune responses during malaria in a controlled environment, CHMI participants are treated relatively early in infection and how our data translate to endemic areas and childhood infections remains to be investigated.

Limitations of our study are the relatively small number of NK cell receptors investigated, including those involved directly in parasite recognition (Sakoguchi et al., 2025; Saito et al., 2017). Additionally, links between CMV, NK cells and parasite control were only investigated in malaria naïve adults in CHMI, and how results translate to children in endemic areas requires further investigation. Due to sample limitations, we were unable to investigate NK cell responses in our complete CHMI cohort of 40 individuals and were unable to test NK cell responsiveness to parasite and/or cytokine stimulation *in vitro* in donors. Furthermore, we couldn’t directly link myeloid compartment changes to NK cell responses in CHMI in the same individuals. Future studies are required to completely dissect the mechanisms underpinning CMV-mediated modulation of the NK cell compartment during malaria.

## Materials and Method

### Ethics

Written informed consent was obtained from all participants. Ethics approval for the use of human samples in the relevant studies was obtained from the Alfred Human Research and Ethics Committee for the Burnet Institute (#288/23 and #166/24), the Human Research and Ethics Committee of the QIMR-Berghofer Medical Research Institute (P1479).

### Study cohorts

Controlled Human Malaria Infection (CHMI) studies were performed as previously described using the Induced Blood Stage Malaria (IBSM) model (Mukhiya et al., 2024; Soon et al., 2025; Chan et al., 2020). Individuals who were malaria naïve were inoculated by intravenous injection of 2,800 *P. falciparum* infected red blood cells and monitored for parasite growth with qPCR (Rockett et al., 2011). Blood samples were collected at baseline (day 0), peak-infection (day 8), day 14/15 (collectively called day 15) and at the end of study (27–36 days post infection, collectively indicated as day 36). Samples were from four studies across six independent infection cohorts, completed between May 2015 and February 2017 (Gaur et al., 2020; McCarthy et al., 2020; Collins et al., 2018). Samples used here were collected from volunteers who consented to donate blood for immunological studies within the parent clinical trial and therefore a sample size estimation was not performed. PBMCs were isolated using Ficoll-Paque (Sigma, USA) density gradient centrifugation and cryopreserved in 10% DMSO/FBS. PBMCs from healthy non-infected individuals were collected by the same processes for analysis of NK responses following *in vitro* stimulation with malaria parasites. For analysis of TLR responses in myeloid cells, whole blood was collected prior to malaria inoculation from additional study participants enrolled in CHMIs ACTRN12621000866808 (Webster et al., 2025) and ACTRN12620000995976 (Barber et al., 2023). For all donors, CMV and EBV seroprevalence was assessed using plasma samples (at day 0 for all CHMI) by commercially available ELISA kits (ab108724 and ab108730), according to manufacturer’s instructions. Clinical score at day 7 and 8 following inoculation was calculated by the number and severity of malaria-related symptoms.

#### Sex as a biological variable

Both male and females were included in this study. For some parent CHMI studies, females of childbearing age were excluded and as such the sex is not evenly distributed.

### *P. falciparum* parasite culture

Packed red blood cells (RBCs) from donors were infected *in vitro* with the *P. falciparum* 3D7 parasite strain (Trager and Jensen, 1976). Packed RBCs for parasite culture were acquired from the Australian Red Cross. *P. falciparum-*infected RBCs (pRBCs) were cultured at 5% haematocrit in Roswell Park Memorial Institute 1640 media (RPMI) supplemented with AlbuMAX II (0.25%) and heat-inactivated human sera (5%). Cultures were incubated at 37 °C in 1% O_2_, 5% CO_2_, and 94% N_2_ gas mixture. Culture media was replaced daily, and parasite stage/parasitemia was monitored by Giemsa-stained blood smears. pRBCs were grown to 15% parasitemia and purified from uninfected RBCs (uRBCs) and early stage pRBCs via magnet separation to enrich mature trophozoite stage pRBCs. Purified pRBCs (>95% purity) were stored at -80 °C following addition of a Glycerolyte cryopreservant.

### Profiling NK cell transcriptional response to *P. falciparum* parasites

PBMCs were thawed in 10% FBS/RPMI and cultured 1:1 cell ratio with 1x10^6^ mature trophozoite stage pRBCs or media (unstimulated control) at 37°C, 5% CO2 in 96-well U-bottom plates for 24 hours. Following culture, NK cells were isolated using negative selection (MACS Miltenyi Biotec, kit#130-092-657) and cell purity and viability quantified by flow cytometry (Supplementary Table S3). Purity and viability were similar between CMV seronegative and seropositive, nor unstimulated compared to parasite stimulated cells (Supplementary Figure 1SA/B). RNA was extracted from NK cells using the QIAGEN PicoPureTM RNA isolation kit (Applied Biosystems™, KIT0204), and RNA quality confirmed with the 2200 TapeStation system (G2964AA) by High Sensitivity RNA ScreenTape (5067- 5579). RNA sequencing libraries were constructed using the NEBNext Single Cell/Low Input RNA Library Prep Kit for Illumina (E6420S) and NEBNext Multiplex Oligos for Illumina (96 Unique Dual Index Primer Pairs) (E6440S). One sample from an unstimulated CMV negative female donor failed RNA extraction and was not sequenced. The libraries were sequenced using a paired-end NextSeq 500/550 high output kit v2.5 (150 cycles) (Cat #20024907).

### *Ex vivo* NK cell phenotyping during CHMI

PBMCs were thawed in 10% FBS/RPMI 1640 and 0.02% Benzonase, and 1x10^6^ cells were stained with antibody LAG3, Fc (1:100, BD Biosciences) and monoblock (1:20, Biolegend) at 37°C, 5% CO2 for 45 min. After two washes (1x PBS), PBMCs were stained with Viadye Red live dead for 15 mins at room temperature (RT), washed twice with 2% FBS/PBS. Then, PBMCs were pre-stained with TCR ψ8 at RT for 15 mins and then surface stained for 15 minutes at RT with fluorescent-tagged antibodies in 2% FBS/PBS (Supplementary Table S4). Following two washes with 2% FBS/PBS, cells were fixed and permeabilized with eBioscience Foxp3/Transcription Factor Staining Buffer for 20 mins on ice. Intracellular staining was performed for 30 mins on ice after 2 washes with 1x perm buffer (Supplementary Table S4) and PBMCs were fixed with BD Stabilizing Fixative (BD Biosciences) and resuspended in 1x PBS. Cells were acquired on the Cytek Aurora 5 laser cytometer within 24 hours.

### TLR whole blood stimulation and intracellular cytokine staining

To assess TLR responsiveness of innate cells 300 µl of whole blood was stimulated with Pam3Csk4/HKLM (TLR1, 0.1 µg/mL; TLR2, 10^8^ cells/mL), LPS (TLR4, 200 ng/mL) or CL075 (TLR7/8, 4 µg/mL), after 1 hour at 37°C Brefeldin A (10 µg/mL) and Monensin (10 µg/mL) were added and stimulated cells incubated for a further 3 hours at 37°C. After 4 hours of stimulation, each sample was resuspended in 420 µL of PROT1-1 (Smart Tube Inc) stabilising fixative according to the manufacturer’s instructions. Cryopreserved whole blood samples were stored at – 80°C. After collection of all samples from each CHMI study, all cryopreserved TLR stimulated whole blood samples were analysed in a single batch. Samples were thawed in an agitating water bath set at room temperature for 10 mins. Thawed samples were transferred into labelled FACS tubes and RBCs lysed according to the manufacturer’s instructions (Smart Tube Inc). In brief, samples were resuspended in 1x Thaw-Lyse buffer, incubated for 10 mins at RT, centrifuged at 600g for 5 minutes at RT. Lysis was repeated, and cell pellets resuspended in 2% FBS/PBS. Resuspended cell pellets were transferred to a 96 V-bottom plate for intracellular cytokine staining. Washed cells were incubated with Fc (1/100, BD Biosciences) and Monocyte Block (1/20, Biolegend) for 10 mins at RT. Cells were resuspended with the surface antibody panel (Supplementary Table S5a/b) and incubated for 15 mins at RT. After washing with 2% FBS/PBS cells were fixed with CytoFix/CytoPerm (BD Biosciences) for 20 mins on ice. After fixation cells were washed with 1 x Permeabilization buffer (BD Biosciences) and resuspended with intracellular antibody mastermix (Supplementary Table S5a/b). Finally, cells were treated with 1x BD Stabilising Fixative (BD Biosciences) for 20 min at RT. Cells were resuspended in 2% FBS/PBS and acquired on a Cytek Aurora 5 within 24 hours.

### Data analysis

#### Flow cytometry

FACS data was acquired using the Cytek Aurora 5 (CA, USA) and manual gating was performed in FlowJo v10 (BD, 2019). For clustering analysis, we used the R package SPECTRE (Ashhurst et al., 2021). Unsupervised clustering was performed and cell clusters were visualized with uniform manifold approximation and projection (UMAP). For functional assays, data was analysed with Flowjo v10 (BD, 2019) and R/R studio was used for statistical analysis and data visualisation.

#### Bulk RNAseq

Raw sequencing reads were first trimmed to remove adapter sequences and low-quality bases using Cutadapt (v1.9). Trimmed reads were then aligned to the Human GRCh37 reference genome, incorporating Ensembl v97 gene models, using STAR (v2.5.2a). Alignment files were processed, sorted, and converted to the required formats using SAMtools (v1.9). Gene and transcript expression levels were quantified with RSEM (v1.2.30), providing normalized expression estimates. Quality assessment of the RNA-seq data was performed using RNA-SeqQC (v1.1.8) to ensure data reliability. All analyses were carried out using Python (v3.6.1) and Perl (v5.22) for scripting and workflow automation. Low-count genes (fewer than 10 counts) were removed, and dispersion was estimated using edgeR (v4.40) workflow in R. A negative binomial distribution via regression models of normalized count data and Wald test was used to compare gene expression variation between paired pre- and post-stimulated samples from CMV negative and CMV positive with the glmmSeq R package (Lewis et al., 2022). The design matrix accounted for random effects of individual samples and included an interaction term between state (pre- vs. post-stimulation) and CMV (CMV negative vs. CMV positive). Fold change due to stimulation was calculated by subtracting the log-transformed response term of unstimulated from stimulated samples. Fold change due to CMV was calculated by subtracting the log-transformed response term of CMV negative individuals from CMV positive individuals for both stimulated and unstimulated samples. We corrected for multiple testing using Storey’s q-value method, defining significance for q-values lower than 0.05 with the q-value R package.

### Statistics

Cellular response comparisons within CMV serostatus was performed using non-parametric testing, Wilcoxon signed rank test. For comparisons between CMV groups the Mann Whitney U test was performed. PMR (calculated as multiplication for a 48 hour period) were calculated by applying a sine-wave growth model to the parasitemia data (Wockner et al., 2020). Speaman’s correlation test was used to assess association between with PMR and cellular responses. P values were not adjusted for comparisons. All statistical analysis were performed in R (version 4.3.2). Graphical outputs were made using the R package ggplot2 (version 3.5.1).

## Data availability

The fastq files and raw counts for transcriptional data generated in this study have been deposited in Gene Expression Omnibus database under accession code GSE278045.

## Code availability

Code used to analyse RNAseq data is available https://github.com/Boyle-Lab-CRDV/Mukhiya_2026_CMV.git

## Acknowledgements

Burnet Institute, University of Melbourne and QIMR-Berghofer acknowledge the traditional custodians of the lands where they are located, the Boonwurrung and Wurundjeri Woi-wurrong people of the Kulin Nation, and the Turrbal and Jagera people.

RBC and human serum were provided by the Australian Red Cross Blood Bank (Melbourne and Brisbane). We thank the participants involved in the CHMI studies and all study clinicians and support staff at QPharm and University of Sunshine Coast. This work was supported by the National Health and Medical Research Council of Australia (program grant 1132975 to J.S.M and C.R.E.); Senior Research Fellowships C.R.E. (1154265), Career Development Award 1141278, Project Grant 1125656, and Ideas Grant 1181932 to MJB, and Ideas grant 2002957 to MJB and BEB); the CSL Centenary Fellowship and the Snow Medical Foundation Fellowship 2022/SF167 to

M.J.B. Burnet Institute is supported by the NHMRC for Independent Research Institutes Infrastructure Support Scheme and the Victorian State Government Operational Infrastructure Support. Medicines for Malaria Venture funded the parent CHMI studies.

## Author contributions

RM, JRL, NK, DA, generated data

RM, JRL, ZP, DO analysed data

JSM, BEB provided clinical samples

JAL, CE, MJB provide supervision

RM, JRL, MJB wrote manuscript with feed-back from all authors

## Notes

*Conflicts of interest statement:* All authors declare no conflicts of interest

### Competing Interest Statement

The authors have declared no competing interest.

