## Supplementary Tables and Figures for "Latent cytomegalovirus disrupts NK cell responses to *P. falciparum* and impairs parasite control"

**Supplementary Table 1: glmmSeq analysis of NK cell response to parasite stimulation in CMV seronegative and positive individuals** (*attached as excel)*

**Supplementary Table 2: Demographic characteristics of CHMI cohort**

| | CMV serostatus | | |
|---|---|---|---|
| | Negative | Positive | P |
| Total, n (%) | 19, (48%) | 21, (52%) | 0.751* |
| Sex, male, n (%) | 18 (95%) | 18 (86%) | 1* |
| EBV, positive, n (%) | 18 (95%) | 17 (81%) | 0.865* |
| Age years, median [IQR] | 26 [20.25-31.75] | 25 [21-29] | 0.86# |

* Chi-square, # wilcox rank sum

**Supplementary Table 3: NK cell panel for purity check:**

| Fluorophore | Marker | Dilution | Cat | Clone | Supplier | Lot |
|---|---|---|---|---|---|---|
| Live dead aqua | Dead | 1:5000 | | | | |
| BV421 | CD56 | 1:100 | 562752 | NCAM16.2 | BD | 1110863 |
| BV650 | CD3 | 1:100 | 317324 | OKT3 | Biolegend | B321523 |
| BV785 | HLADR | 1:100 | 307642 | L243 | Biolegend | B324516 |
| FITC | VD1 | 1:50 | TCR2730 | TS8.2 | Invitrogen | UB277210 |
| PerCp-Cy5.5 | CD19 | 1:25 | 561295 | HIB19 | BD | 1159287 |
| PE | CD14 | 1:50 | 561707 | M5E2 | BD | 1349562 |
| PE-Daz | CD64 | 1:100 | 305032 | 10.1 | Biolegend | B292051 |
| APC | VD2 | 1:50 | 331418 | B6 | Biolegend | B352327 |
| AF700 | CD16 | 1:100 | 302026 | 3G8 | Biolegend | B339559 |

**Supplementary Table 4: NK cell *ex vivo* phenotyping panel in CHMI:**

| Fluorophore | Marker | Dilution | Cat | Clone | Supplier | Lot |
|---|---|---|---|---|---|---|
| ViadyeRed | Live dead | 1:1000 | | | Cytek | |
| PerCPCy5.5 | TCR γδ | 1:10 | 564157 | B1 | BD | 2207690 |
| BUV661 | LAG3 | 1:25 | 376-2239-42 | 3DS223H | Invitrogen | 2783561 |
| BUV737 | CD56 | 1:100 | 612766 | NCAM16.2 | BD | 1210146 |
| BUV805 | CD3 | 1:400 | 612893 | SK7 | BD | 3212687 |
| BV480 | CD7 | 1:400 | 566119 | MT701 | BD | 2059133 |
| BV570 | HLA-DR | 1:66 | 307638 | I243 | Biolegend | B382459 |
| BV650 | CD14 | 1:200 | 301836 | M5E2 | Biolegend | B360116 |
| BV711 | NKp30 | 1:50 | 563383 | P30-15 | BD | 2131202 |
| BV786 | PD1 | 1:50 | 563789 | EH12.1 | BD | 3276824 |
| BB515 | CD86 | 1:400 | 564545 | 2331 | BD | 9261900 |
| PE | NKG2C | 1:400 | 375004 | S19005E | Biolegend | B370170 |
| PE-CY7 | NKG2A | 1:100 | 375114 | S19004C | Biolegend | B401967 |
| APC | TCR Vδ2 | 1:100 | 331418 | B6 | Biolegend | B352327 |
| efluor660 | CD57 | 1:100 | 50-0577-42 | TB01 | Invitrogen | 2518387 |
| APCfire810 | CD38 | 1:200 | 303550 | HIT2 | Biolegend | B394848 |
| BV421 | Perforin | 1:1600 | 563393 | DG9 | BD | 3254954 |
| AF488 | Granulysin | 1:16 | 558254 | RB1 | BD | 0325603, |
| PE-Fire810 | TIGIT | 1:25 | 372745 | A15153G | Biolegend | B409947 |
| Pe-dazzle594 | CD85j | 1:33 | 333716 | GHI/75 | Biolegend | B375814 |
| AF700 | CD16 | 1:100 | 302026 | 3G8 | Biolegend | B384538 |
| Percp-eFlour710 | CD19 | 1:50 | 46-0198-42 | SJ25C1 | Invitrogen | 2005232 |
| APC | GranzB | 1:1600 | 372204 | QA16A02 | Biolegend | B281725 |

**Supplementary Table 5a: Innate cell whole blood ICS panel**

| Fluorophore | Marker | Dilution | Cat | Clone | Supplier | Lot |
|---|---|---|---|---|---|---|
| *Surface stain* | | | | | | |
| BUV496 | CD19 | 1:100 | 12938 | SJ25C1 | BD | 1182272 |
| BUV737 | CD64 | 1:50 | 612776 | 10.1 | BD | 3018874 |
| BUV805 | CD14 | 1:50 | 612902 | M5E2 | BD | 2213039 |
| BV480 | CD86 | 1:50 | 566131 | 2331 | BD | 2136488 |
| BV510 | CD1c | 1:100 | 331534 | L161 | BioLegend | B281571 |
| BV570 | CD33 | 1:50 | 303417 | WM53 | BioLegend | B378792 |
| BV605 | CD303 | 1:50 | 3542274 | 201A | BioLegend | B382651 |
| BV605 | CD123 | 1:100 | 306026 | 6H6 | BioLegend | B379483 |
| BV650 | CD11c | 1:200 | 337238 | Bu15 | BioLegend | B323827 |
| BV750 | CD56 | 1:33 | 362556 | 5.1H11 | BioLegend | B363758 |
| BV785 | HLA-DR | 1:100 | 307642 | L243 | BioLegend | B324516 |
| BB515 | CD3 | 1:100 | 565100 | UCHT1 | BD | 1187749 |
| PerCPCy5.5 | CCR2 | 1:50 | 357204 | K036C2 | BioLegend | B328029 |
| PE-FIRE640 | CD66b | 1:200 | 392918 | 6/40c | BioLegend | B348028 |
| AF700 | CD16 | 1:50 | 302026 | 3G8 | BioLegend | B384538 |
| *Intracellular stain* | | | | | | |
| BUV395 | TNF | 1:25 | 563996 | MAb11 | BD | 1271128 |
| BV421 | IL-12 | 1:25 | 565023 | C8.6 | BD | 2182102 |
| FITC | IL-1b | 1:25 | 11-7013-42 | CRM56 | Thermofisher Invitrogen | 2608896 |
| PE | IL-10 | 1:10 | 559337 | JES3-9D7 | BD | 307185 |
| PE-Cy7 | IL-6 | 1:100 | 501120 | MQ2-13A5 | BioLegend | B320181 |
| AF647 | MCP-1 | 1:50 | 563496 | 5D3-F7 | BD | 9338878 |
| APC | IFNa | 1:5 | 130-092-602 | LT27:295 | Miltenyi Biotec | 5230310707 |

**Supplementary Table 5b: Innate cell whole blood ICS panel**

| Fluorophore | Marker | Dilution | Cat | Clone | Supplier | Lot |
|---|---|---|---|---|---|---|
| *Surface stain* | | | | | | |
| BUV496 | CD19 | 1:100 | 612938 | SJ25C1 | BD | 1182272 |
| BUV737 | CD64 | 1:50 | 564425 | 10.1 | BD | 8200909 |
| BUV805 | CD14 | 1:50 | 612902 | M5E2 | BD | 1092492 |
| BV480 | CD86 | 1:50 | 566131 | 2331 | BD | 9064680 |
| BB515 | CD3 | 1:100 | 564466 | UCHT1 | BD | 1187749 |
| BV510 | CD56 | 1:100 | 318340 | HCD56 | Biolegend | B367784 |
| BV570 | CD33 | 1:50 | 303417 | WM53 | Biolegend | B332083 |
| BV605 | CD123 | 1:100 | 306026 | 6H6 | Biolegend | B265668 |
| BV650 | CD11c | 1:200 | 117310 | B915 | Biolegend | B323827 |
| BV785 | HLADR | 1:100 | 307642 | L243 | Biolegend | B324516 |
| PerCPCy5.5 | CCR2 | 1:50 | 357204 | K036C2 | Biolegend | B307717 |
| BV605 | CD303 | 1:50 | 354224 | 201A | Biolegend | B349636 |
| PE-FIRE640 | CD66b | 1:100 | 392918 | 6140c | Biolegend | B321127 |
| AF700 | CD16 | 1:100 | 302026 | 3C78 | Biolegend | B333714 |
| APC-FIRE 750 | CD1c | 1:33 | 331545 | L161 | Biolegend | B336699 |
| *Intracellular stain* | | | | | | |
| AF647 | MCP1 | 1:50 | 563496 | 503-F7 | BD | 9338878 |
| BV750 | TNF | 1:5 | 566359 | Mab11 | BD | 1133444 |
| BV421 | IL12 | 1:25 | 565023 | C8-6 | BD | 7263688 |
| PE-Cy7 | IL6 | 1:100 | 501119 | MQ2-13A5 | Biolegend | B281369 |
| FITC | IL1b | 1:25 | 11-7018-42 | CRM56 | Invitrogen | 2527372 |

### Supplementary Figures

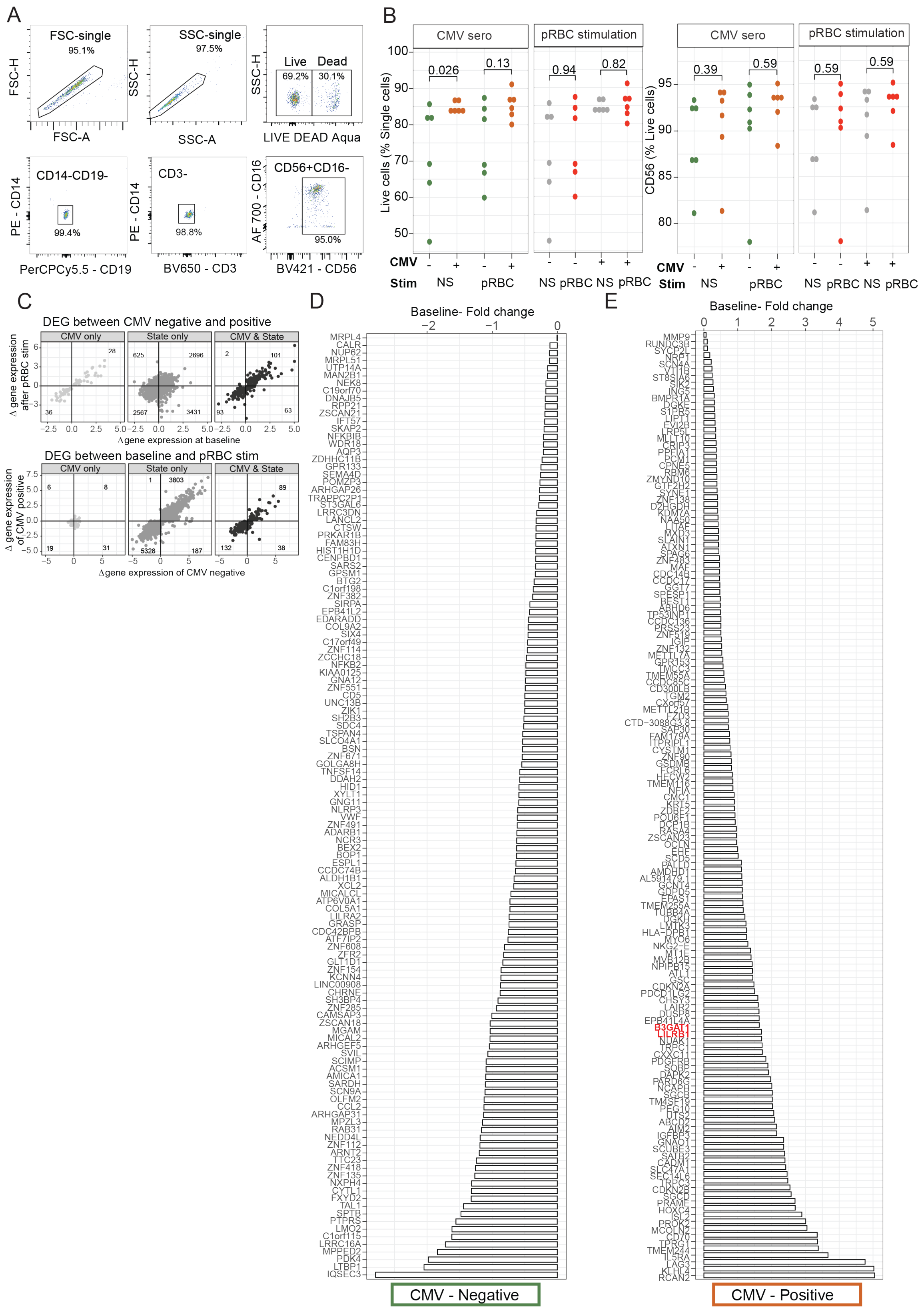

***Supplementary Figure 1: Transcriptional analysis of NK cells in response to in vitro parasite stimulation between CMV seronegative and seropositive individuals***

***(A)*** *Gating example of post isolation QC flowcytometry of NK cells in unstimulated.* ***(B)*** *Percentage of live and CD56 cells assessed via flow cytometry after cell-sorting.* ***(C)*** *Scatter plots of expression of DEGs grouped based on if they were significant for "CMV", "State" or "CMV/State Interaction". Left plots show gene expression at baseline compared to after stimulation with pRBCs and right plots show gene expression levels in CMV seronegative compared to CMV seropositive individuals.* ***(D)*** *DEGs identified in glmmSeq that are relatively higher in CMV sero-negative (left panel) and CMV positive (right panel) in unstimulated cells. Related to Figure 1.*

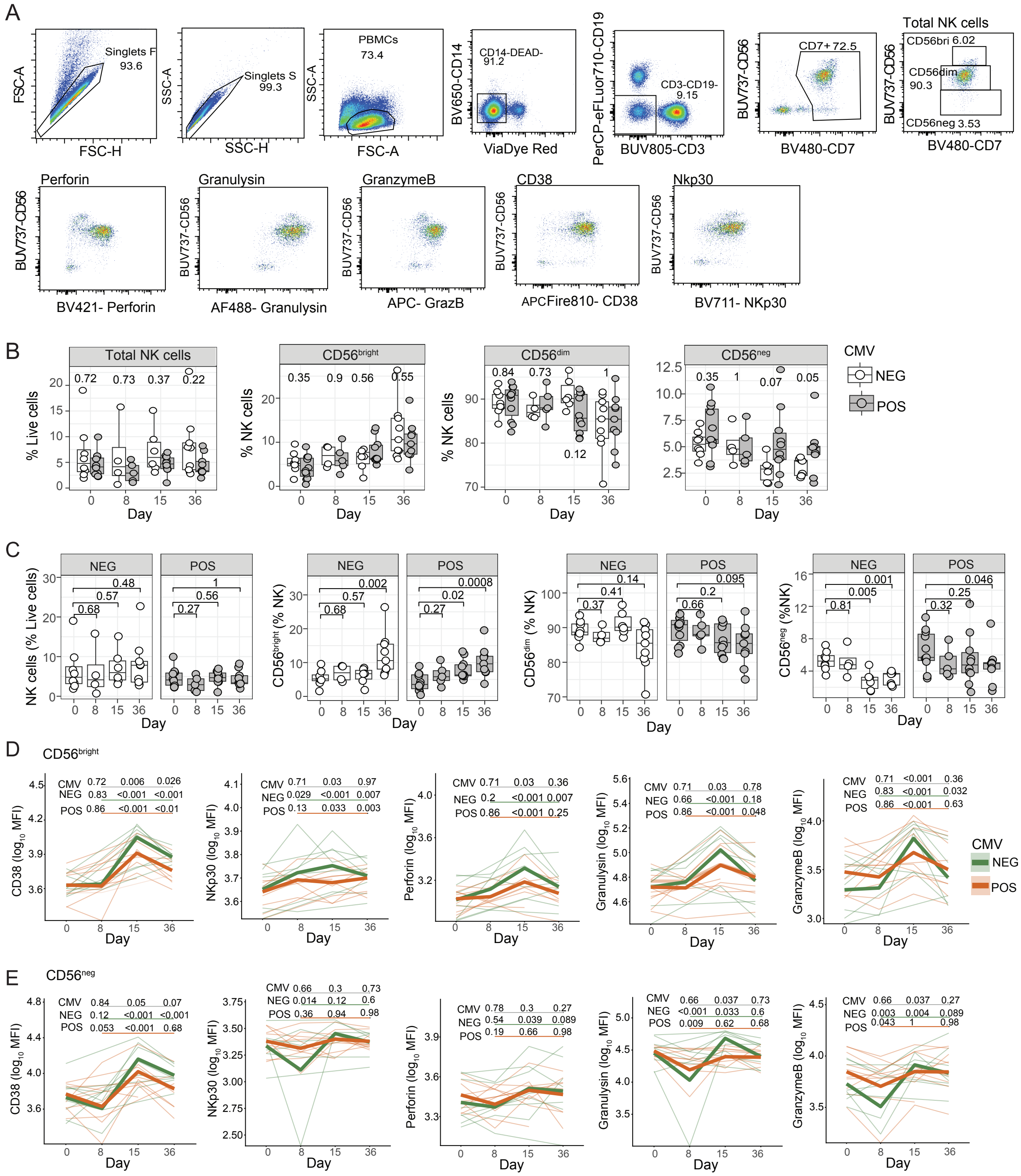


***Supplementary Figure 2: NK cells activation in CHMI in CMV seronegative and CMV seropositive individuals***

*NK cells were analysed during CHMI in CMV seronegative (n=8) and CMV seropositive individuals (n=9). (**A**) Gating figure of NK cells in CHMI. (**B**) Proportion of NK subsets at Day*

*0. **(C)** Total NK cells and its subsets proportions across CHMI. **(D)** Total NK cells and its subsets proportion during malaria in CMV negative and positive separately. **(E)** MFI of activation markers such as Perforin, GrzB, Granulysin, NKp30 and CD38 in CD56bri during malaria. **(F)** MFI of activation markers such as Perforin, GrzB, Granulysin, NKp30 and CD38 in CD56neg during malaria. For **E/F**, data are $log_{10}$ MFIs of markers with thin lines representing individual data coloured by CMV serostatus, and bold lines representing the mean of the predicted values from the fitted models for each group. P values are from linear mixed effect models. CMV is p values for the interaction term between each timepoint (compared to day 0) and CMV serostatus (underlined in grey). NEG/POS are P values for the comparison between day 0 and each subsequent timepoint for CMV seronegative individuals (NEG, underlined in green) and CMV seropositive individuals (POS, underlined in orange) which were determined from contrasts. See also Supplementary Figure S2.*

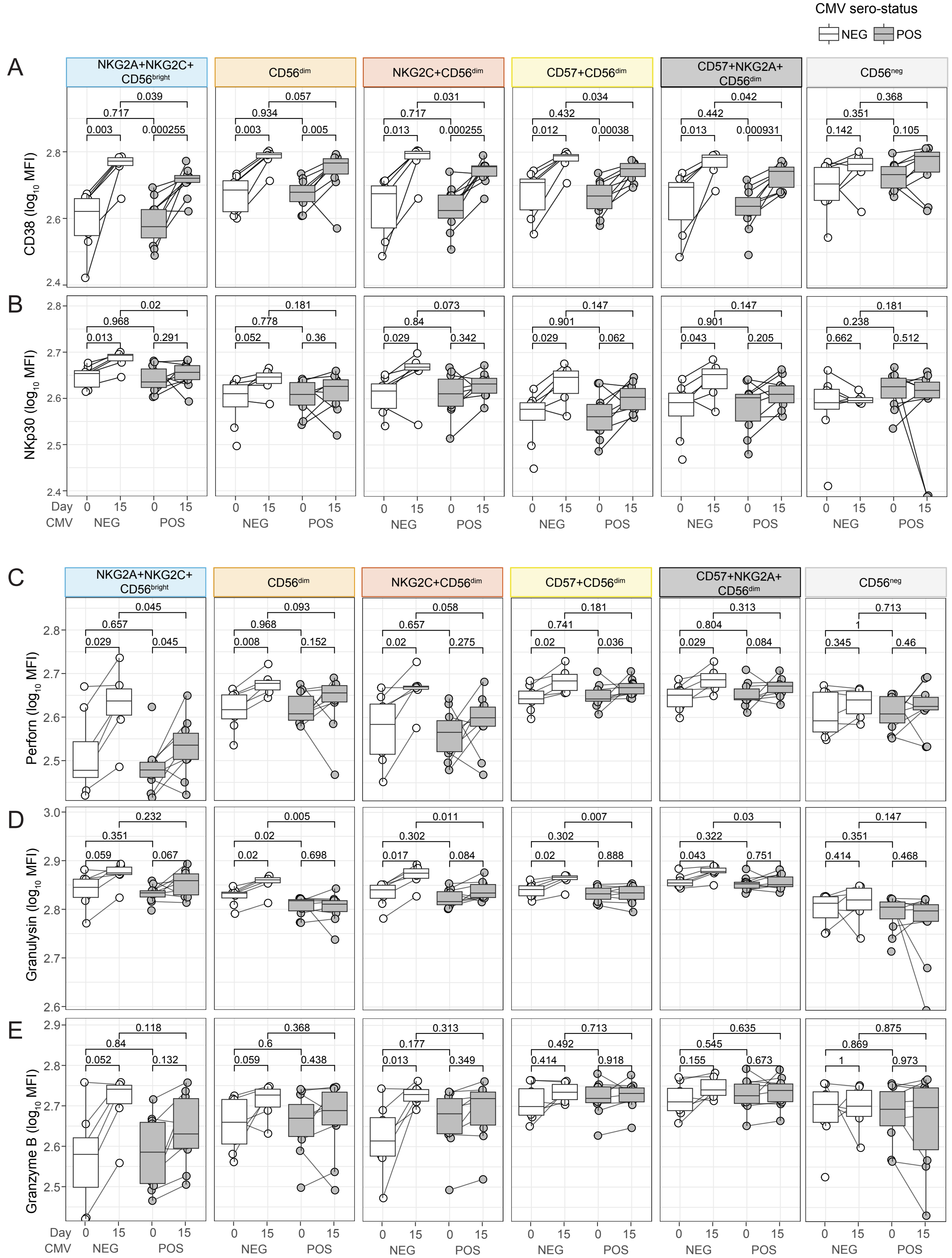

CMV sero-status
NEG
POS
A
B
C
D
E
NKG2A+NKG2C+ CD56$^{bright}$
CD56$^{dim}$
NKG2C+CD56$^{dim}$
CD57+CD56$^{dim}$
CD57+NKG2A+ CD56$^{dim}$
CD56$^{neg}$
CD38 ($\log_{10}$ MFI)
NKp30 ($\log_{10}$ MFI)
Perforn ($\log_{10}$ MFI)
Granulysin ($\log_{10}$ MFI)
Granzyme B ($\log_{10}$ MFI)
Day
CMV
0 15 0 15
NEG POS
0.039 0.717 0.003 0.000255
0.057 0.934 0.003 0.005
0.031 0.717 0.013 0.000255
0.034 0.432 0.012 0.00038
0.042 0.442 0.013 0.000931
0.368 0.351 0.142 0.105
0.02 0.968 0.013 0.291
0.181 0.778 0.052 0.36
0.073 0.84 0.029 0.342
0.147 0.901 0.029 0.062
0.147 0.901 0.043 0.205
0.181 0.238 0.662 0.512
0.045 0.657 0.029 0.045
0.093 0.968 0.008 0.152
0.058 0.657 0.02 0.275
0.181 0.741 0.02 0.036
0.313 0.804 0.029 0.084
0.713 1 0.345 0.46
0.232 0.351 0.059 0.067
0.005 0.02 0.02 0.698
0.011 0.302 0.017 0.084
0.007 0.302 0.02 0.888
0.03 0.322 0.043 0.751
0.147 0.351 0.414 0.468
0.118 0.84 0.052 0.132
0.368 0.6 0.059 0.438
0.313 0.177 0.013 0.349
0.713 0.492 0.414 0.918
0.635 0.545 0.155 0.673
0.875 0.869 1 0.973

***Supplementary Figure 3: Activation and cytotoxicity in NK cell subsets during CHMI.***

*Activation (**A** - CD38 and **B** - NKp30) and cytotoxic (**C** - Perforin, **D** - granulysin, **E** – Granzyme B) were quantified (median fluorescence intensity, MFI) in NK cell subsets that were not otherwise modulated by CMV infection. Expression was compared between day 0 and day 16 in CMV seronegative (n = 8) and CMV seropositive individuals (n = 9), and expression at day 0 or at day 16 was compared between groups. Data are Tukey boxplots with the median, $25^{th}$ and $75^{th}$ percentiles. The upper and lower hinges extend to the largest and smallest values, respectively but not further than 1.5X IQR from the hinge. Individual data are shown as points. For comparisons between groups p is Mann-Whitney U test. For comparisons within groups between days p is Wilcoxon signed-rank test.*

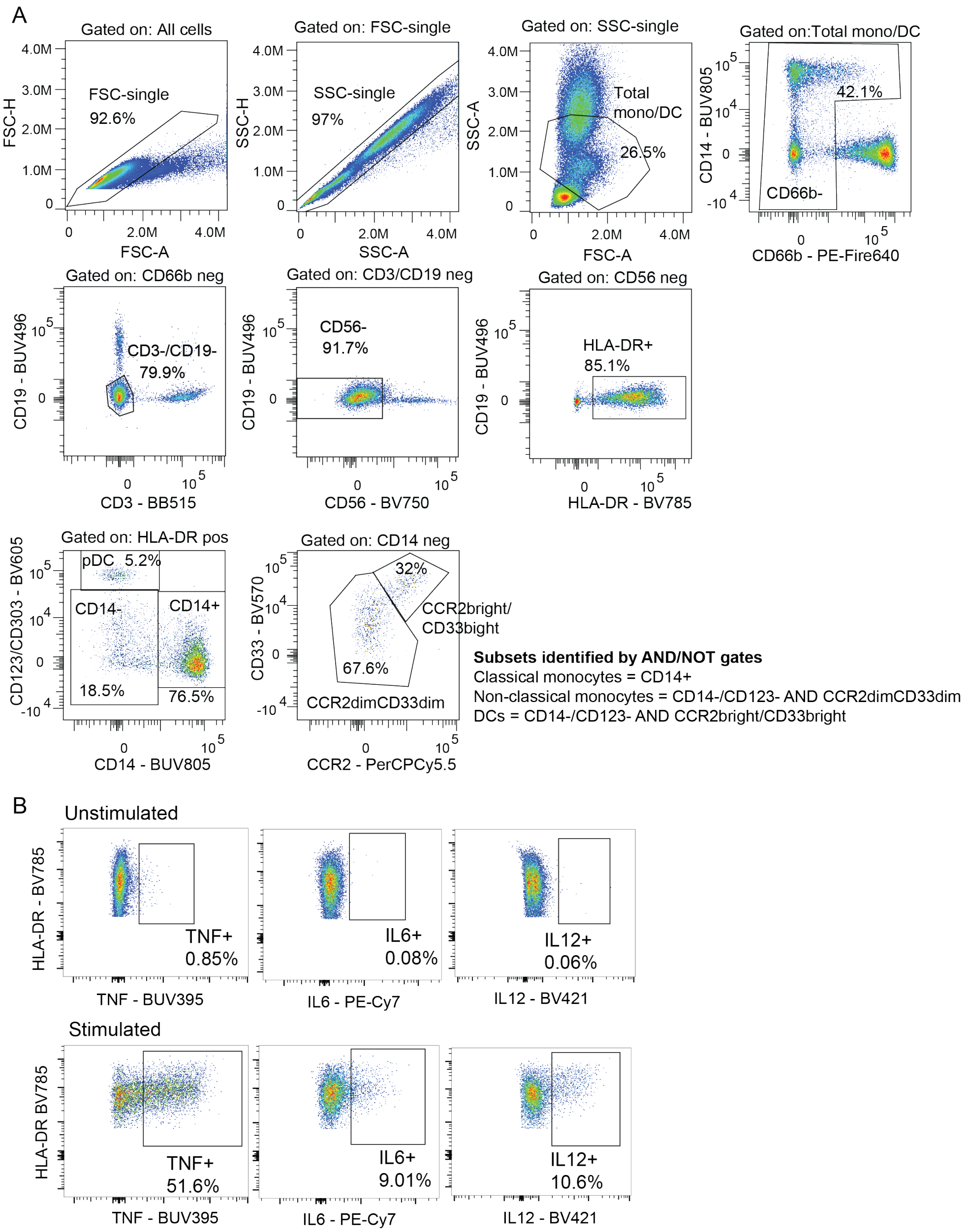


***Supplementary Figure 4: Cytokine response following TLR stimulation of innate cells. (A)*** *Whole blood flow cytometry gating example for identifying classical, non-classical monocytes, DCs and pDCs after stimulation. Total myeloid cells were identified as CD66b/CD3/CD19/CD56 neg HLADR+, classical monocytes were identified as CD14+, non-classical monocytes (were identified as CD14-/CCR2dim/CD33dim, DCs identified as CD14-/CCR2bright/CD33bright and pDCs as CD123/CD303+.* ***(B)*** *Example flow cytometry plots of cytokine production from total myeloid cells, unstimulated (top line) and stimulated (TLR4) (bottom line).*
